# *In silico* Repositioning of Approved Drugs Against *Schistosoma mansoni* Energy Metabolism Targets

**DOI:** 10.1101/397059

**Authors:** Nicole Melo Calixto, Daniela Braz dos Santos, José Clecildo Barreto Bezerra, Lourival de Almeida Silva

**Affiliations:** Department of Bioinformatics, Federal Institute of Education, Science and Technology Goiano -Campus Ceres, Ceres, Goiás, Brazil.; LAERPH- Laboratory of Parasite-Host Relationship Study, Institute of Tropical Pathology and ublic Health of the Federal University of Goiás, Goiânia, Goiás, Brazil.

**Keywords:** Schistosomiasis, Chemogenomics, Drug Discovery, Drug Repositioning

## Abstract

Schistosomiasis is a neglected parasitosis caused by *Schistosoma* spp. Praziquantel is used for the chemoprophylaxis and treatment of this disease. Although this monotherapy is effective, the risk of resistance and its low efficiency against immature worms compromises its effectiveness. Therefore, it is necessary to develop new schistosomicide drugs. However, the development of new drugs is a long and expensive process. The repositioning of approved drugs has been proposed as a quick, cheap, and effective alternative to solve this problem. This study employs chemogenomic analysis with use of bioinformatics tools to search, identify, and analyze data on approved drugs with the potential to inhibit *Schistosoma mansoni* energy metabolism enzymes. The TDR Targets Database, Gene DB, Protein, DrugBank, Therapeutic Targets Database (TTD), Promiscuous, and PubMed databases were used. Fifty-nine target proteins were identified, of which 18 had one or more approved drugs. The results identified 20 potential drugs for schistosomiasis treatment, all approved for use in humans.

## Introduction

Schistosomiasis is a neglected parasitic illness caused by *Schistosoma* trematodes, namely three species of medical importance: *S. mansoni*, *S. haematobium*, and *S. japonicum*. In 2016, transmission of this illness was recorded in 78 countries, with about 206 million people requiring preventive chemotherapy. However, approximately 88 million people received treatment in 52 countries with a high or moderate prevalence of this parasitosis [1].

For schistosomiasis treatment and chemophylaxis, the World Health Organization (WHO) recommends the use of the drug praziquantel (PZQ) [2], because it is an anthelmintic that is effective in a single oral dose, contains low toxicity, and has a relatively low cost [3,4]. PZQ acts as an antagonist of calcium ion channels (Ca^2+^) that induce the influx of Ca^2+^, resulting in muscular spasms and paralysis of worms in the adult stage [4,5]. Although this drug is effective, the appearance of strains resistant to PZQ, as well as its low effectiveness against immature worms, make research and new drug development extremely necessary [6–8].

For this reason, the investigation of target molecules that act in the metabolism of *Schistosoma* spp. and have potential to interact with compound candidates for anti-schistosomiasis therapy is necessary [9,10]. This strategy focuses on energy metabolism enzymes, as they are considered interesting targets for inhibitor development because they reduce energy production capacity, consequently contributing to the death and elimination of the parasite [11].

Conventional research and development strategies for novel drugs are considered high-cost (estimated at millions of dollars), time-consuming (between 10–17 years), and risky. The serious side-effects and diminished efficacy in humans that can occur during clinical trials are common factors leading to a compound not being approved for commercialization [12,13]. Moreover, pharmaceutical industries are not interested in investing in the development and production of new drugs for treating Neglected Tropical Diseases (NTDs), because they offer a low profit return [14].

To overcome these difficulties, the repositioning of approved drugs is proposed, which offers an excellent cost-benefit ratio by diminishing the time (between 3–12 years) and money spent compared to the time and cost of developing new compounds. This strategy consists of identifying new therapeutic indications for already approved drugs to treat other illnesses and/or those with already discovered targets [10,13,15–18]. *In silico* analysis, guided by databases that provide information about potential off-target drug interactions, allows the prediction of side effects and the identification of compounds that are candidates for repositioning [19–21]. Using this strategy, the probability of success increases during *in vitro* and *in vivo* tests, the risk of failure is reduced, and the experimentation on animals is rationalized [15,22].

In this study, the following open access databases were used: TDR Targets Database, Gene DB, Protein, Therapeutic Targets Database (TTD), DrugBank, Promiscuous, and PubMed. These databases complement each other to provide information on the profiles of hundreds of molecular targets and approved drugs, including data about pharmacology, toxicity, pharmacogenomics, and clinical screenings. Using this information, the identification of *S. mansoni* energy metabolism-inhibiting enzymes was possible, to enable the *in silico* repositioning of approved drugs for use in anti-schistosomiasis therapy.

## Materials and methods

The methodology utilized is an adaptation of the methodology used by Bispo et al. [23] and Silva et al. [24].

### Identification of *Schistosoma mansoni* energy metabolism targets

The initial target screening was performed using the TDR Targets Database version 5.0 (Fig 1a) (http://tdrtargets.org/) [25]. On the initial page, the “Targets” option was selected. Next, in the “Select pathogen species of interest” field, the *Schistosoma mansoni* option was selected as the species of interest. The filters field contained the following information: Name/Annotation, Features, Structures, Expression, Antigenicity, Phylogenetic distribution, Essentiality, Validation data, Druggability, Assayability, and Bibliographic references. The number of filters was limited to increase the chances of getting results. For this reason, the only expanded filter was “Name/Annotation,” and the “Energy metabolism” option was selected in the “KEGG high-level pathway” field. Finally, the “Search” option was selected. The database returned a table containing the identity (ID) of each gene of interest.

**Fig 1.**
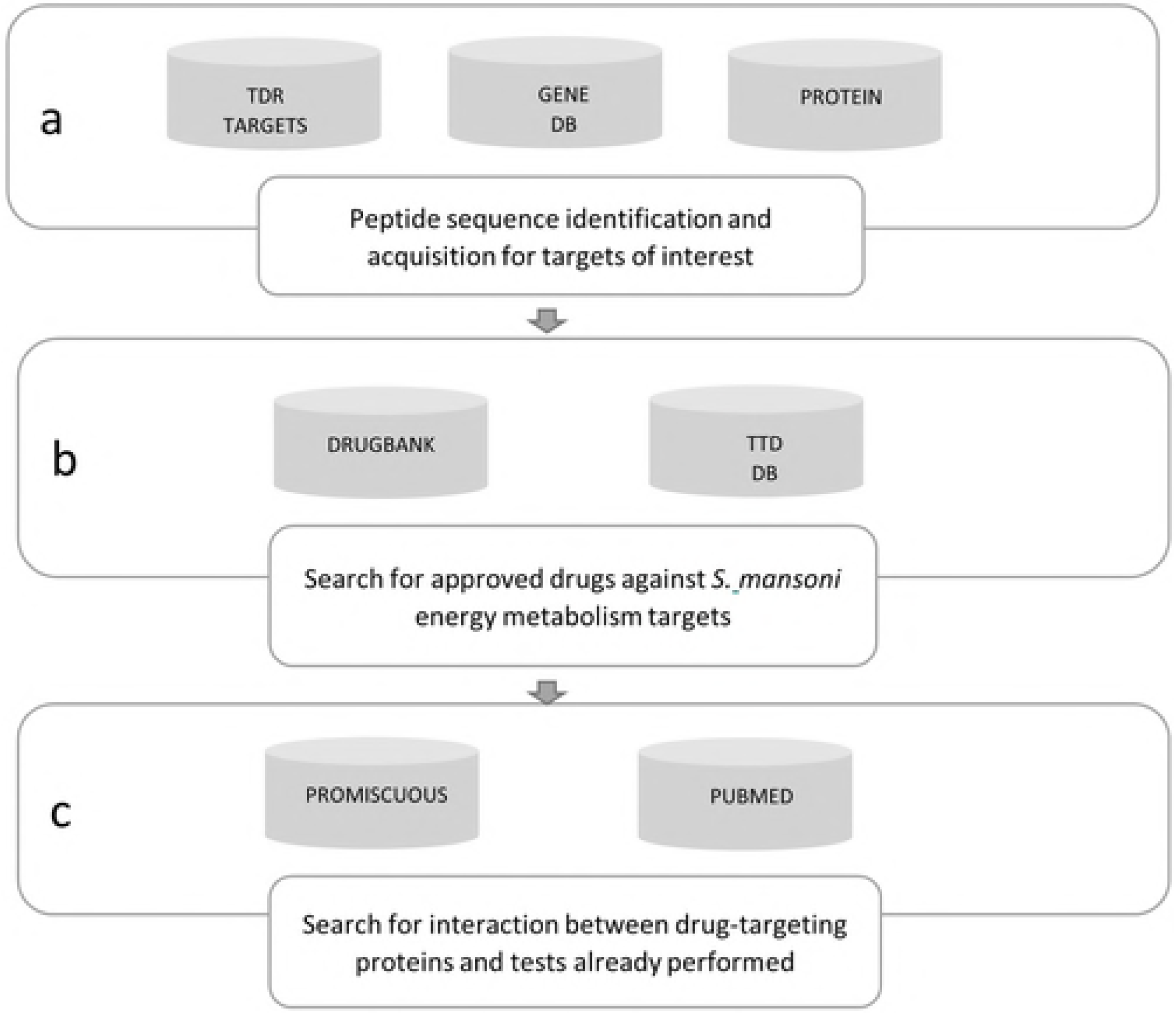
Research pipeline used to identify potential anti-schistosomiasis drugs. a) TDR Targets Database supplied the *S. mansoni* energy metabolism target identity; Gene DB and Protein were used to obtain each target’s peptide sequence. b) DrugBank and TTD were used to search for approved drugs. c) Promiscuous was used to identify the interactions between the drug-target protein and possible side effects of the interaction; PubMed was used to acquire articles on *in vitro* and/or *in vivo* tests with the identified drugs.

After the genes of interest list was produced, the ID of each gene was used to obtain the targets’ peptide sequences form the Gene DB version 2.5 database (Fig 1a) (http://www.genedb.org/) [26]. On the initial Gene DB page, each gene id was inserted into the “Name/Product” field and, right after, “*S. mansoni*” was selected in the “All organism” field, and the search was executed. The Protein database (http://www.ncbi.nlm.nih.gov/protein/) was employed when the gene of interest did not appear in the Gene DB search results. On the initial Protein page, the gene ids were inserted in the “Gene name” field, followed by *Schistosoma mansoni* in the “Organism” field. The gene ids and their respective peptides were tabulated in an Excel spreadsheet.

Peptide sequences were obtained from the DrugBank (http://www.drugbank.ca/) and TTD (http://bidd.nus.edu.sg/group/cjttd/) databases, and this provided basic information about hypothetical drug targets and their corresponding inhibitors.

### Search for approved drugs for the selected targets: general strategy in DrugBank and TTD

DrugBank version 4.0 (Fig 1b) stores information on drugs and their targets. This database contains approximately 8,000 compounds, including 1,700 Food and Drug Administration (FDA)-approved drugs and 6,000 experimental drugs [27].

Another database utilized was the TTD update 2016 (Fig 1b) that provides information on drug targets, such as sequence, function, and three-dimensional structure. At the time of the search, TTD contained 2,589 targets and around 31,614 drugs, of which 2,071 were approved and 7,291 were in the clinical trial phase [28].

In these two databases, the general strategy was based on the principle of target similarity, where each *S. mansoni* energy metabolism target was compared with other targets that already have approved drugs for treatment. All homologous drug targets with an expectation value (*E-value*) less than 1 × 10^-5^ obtained from the databases were input into a table as potential targets.

### Specific commands for the DrugBank database

On the initial DrugBank page, in the “Search” option, “Target Sequences” was selected. Then, we inserted the protein sequence for consultation. The *BLAST* parameters offered by the database were maintained and in the "drug types" filter, the option "approved” was marked.

### Specific commands for the Therapeutic Targets Database

In the initial TTD page, the “Target Similarity” option was selected and the peptide sequence was introduced into the “Input your protein sequence in FASTA format” field, and the “Search” function was executed. All the results with an E-value less than 1 × 10^-5^ were considered for analysis.

### Criteria for the compilation of expected targets

From the results obtained regarding potential drug targets, we select those that interacted with compounds approved for clinical use in human beings. The following parameters were introduced in the table for targets with positive outcomes: “Name(s) of homologous target(s)” (Drug Bank and TTD), “Target ID(s)" (DrugBank and TTD), “Pharmacological indication” (DrugBank), and “Drug approval phase” (DrugBank and TTD) (Table 1).

**Table 1.** Identified targets and their function in *S. mansoni* energy metabolism.

| Number | Target(s) | ID(s)<sup>a</sup> | Function |
| --- | --- | --- | --- |
| 02 | Cytochrome - C oxidase | *Smp_128610*<br>*Smp_144000* | Reduction of O<sub>2</sub> to H<sub>2</sub>O is supplemented by the extrusion of four protons in the intramitochondrial compartment |
| 01 | Glutamate synthase | *Smp_128380.2* | Chemical reactions and pathways that result in glutamate formation, anion of 2-amino pentatonic acid |
| 01 | Alcohol dehydrogenase | *Smp_044440* | Involved in the oxidation process of ethanol for conversion into acetyl-CoA |
| 03 | ATP Synthase | *Smp_029390*<br>*Smp_082120*<br>*Smp_038100* | Involved in the transport of protons connected to ATP synthesis |
| 01 | NADH-ubiquinone oxidoreductase | *Smp_038870.1* | Acts as an NADPH oxireductor |
| 01 | Aminomethyltransferase | *Smp_128980* | Acts on glycine decarboxylation through the cleavage system |
| 01 | Succinate dehydrogenase | *Smp_061230* | Involved in the tricarboxylic acid cycle |
| 02 | Glutaminase | *Smp_147040*<br>*Smp_091460* | Involved in the glutamine metabolic process |

<sup>a</sup>ID – Target identity

### Search for interaction between drugs identified with other protein targets and the possible side effects of this relationship

Promiscuous version 3.0 (http://bioinformatics.charite.de/promiscuous/) contains information on 25,000 drugs, 1,100 different side effects, and 12,000 protein targets. Moreover, this database makes information available on protein-protein and drug-protein interactions [29].

Promiscuous offers different search strategies to the user: drug name, target by interest, and metabolic pathway. In the present study, the search strategy chosen was to use the drug name. On the initial page, the “Drugs” option was selected, and each drug name was inserted into the "Drug name" field. Subsequently, the code of each drug was inserted into the PubChem field, found at https://pubchem.ncbi.nlm.nih.go [30], and the “Only with targets” option was selected. Information on the interaction between the drug and protein target and the eventual side effects were separately included in a table (Table 2).

**Table 2.**
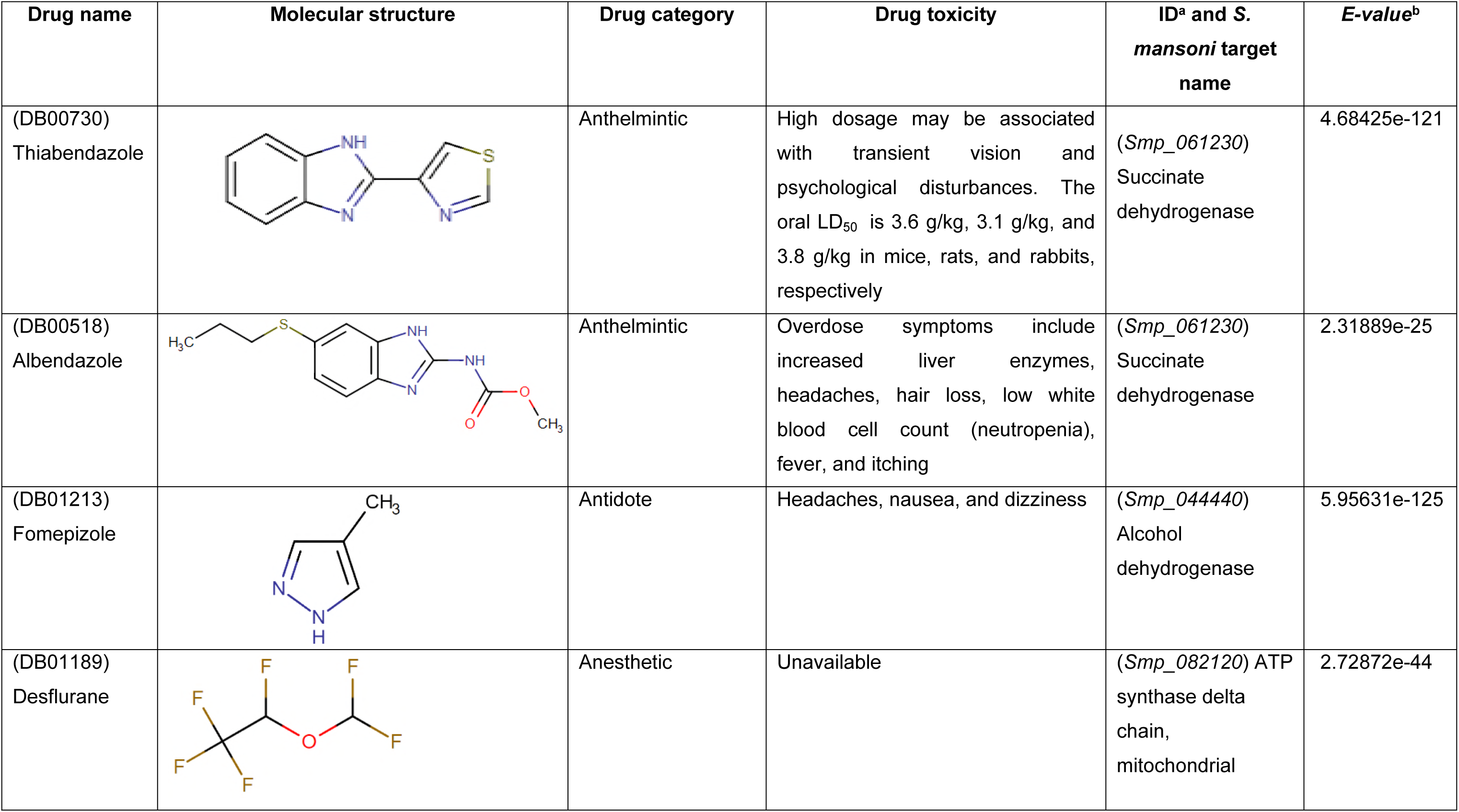
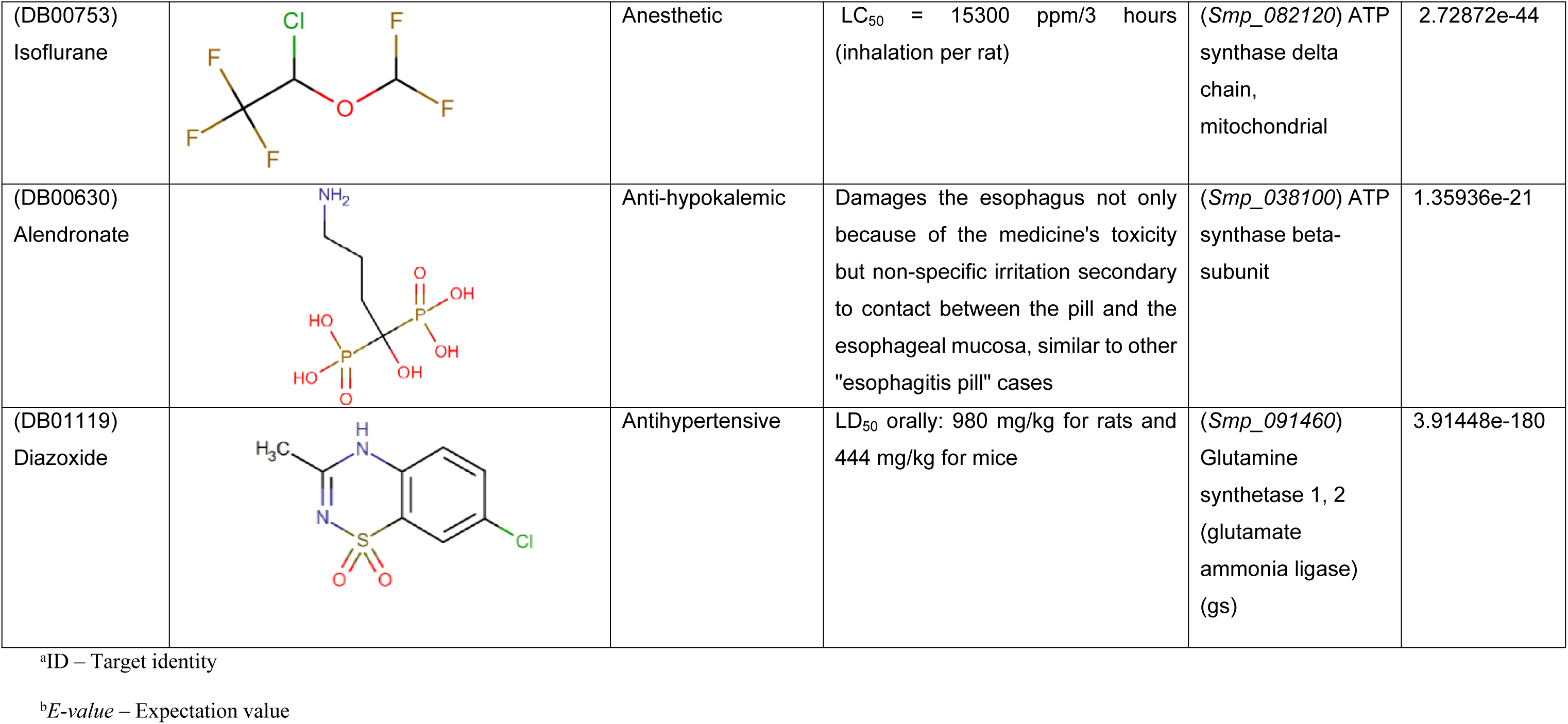
The new associations between *Schistosoma mansoni* energy metabolism targets and drugs identified in the present study.

| Drug name | Molecular structure | Drug category | Drug toxicity | ID<sup>a</sup> and *S. mansoni* target name | *E*-value<sup>b</sup> |
| --- | --- | --- | --- | --- | --- |
| (DB00730)<br>Thiabendazole |  | Anthelmintic | High dosage may be associated with transient vision and psychological disturbances. The oral LD<sub>50</sub> is 3.6 g/kg, 3.1 g/kg, and 3.8 g/kg in mice, rats, and rabbits, respectively | (*Smp_061230*)<br>Succinate dehydrogenase | 4.68425e-121 |
| (DB00518)<br>Albendazole |  | Anthelmintic | Overdose symptoms include increased liver enzymes, headaches, hair loss, low white blood cell count (neutropenia), fever, and itching | (*Smp_061230*)<br>Succinate dehydrogenase | 2.31889e-25 |
| (DB01213)<br>Fomepizole |  | Antidote | Headaches, nausea, and dizziness | (*Smp_044440*)<br>Alcohol dehydrogenase | 5.95631e-125 |
| (DB01189)<br>Desflurane |  | Anesthetic | Unavailable | (*Smp_082120*) ATP synthase delta chain, mitochondrial | 2.72872e-44 |
| (DB00753)<br>Isoflurane |  | Anesthetic | LC<sub>50</sub> = 15300 ppm/3 hours<br>(inhalation per rat) | (*Smp_082120*) ATP<br>synthase delta<br>chain,<br>mitochondrial | 2.72872e-44 |
| (DB00630)<br>Alendronate |  | Anti-hypokalemic | Damages the esophagus not only because of the medicine's toxicity but non-specific irritation secondary to contact between the pill and the esophageal mucosa, similar to other "esophagitis pill" cases | (*Smp_038100*) ATP<br>synthase beta-subunit | 1.35936e-21 |
| (DB01119)<br>Diazoxide |  | Antihypertensive | LD<sub>50</sub> orally: 980 mg/kg for rats and 444 mg/kg for mice | (*Smp_091460*)<br>Glutamine<br>synthetase 1, 2<br>(glutamate<br>ammonia ligase)<br>(gs) | 3.91448e-180 |

<sup>a</sup>ID – Target identity
<sup>b</sup>*E-value* – Expectation value

After obtaining results of interactions between drugs and protein targets and their side effects, the “Show interactive network” option was selected, with the objective of integrating the results in graphs on the interactive network and representing the relationship between cited compounds. The search result was saved in the export network to XGMML section, and the archive was exported to Cytoscape software version 2.0 (http://cytoscape.org/), making it possible to visualize the interaction networks [31].

### List of drugs already tested against *Schistosoma* spp

Lastly, the PubMed database (http://www.ncbi.nlm.nih.gov/pubmed) was used to verify if the identified drugs had already been tested against *Schistosoma spp*. The definition of “test” in this study included *in vitro* and/or *in vivo* tests against the three species of medical importance: *S. mansoni*, *S. haematobium*, and *S. japonicum*. Drugs were classified as “not tested” when no related publication could be found. In those cases, one of the following search terms was introduced into PubMed: (“drug name” [MeSH Terms] OR “drug name” [All Fields]) AND (“Schistosomiasis” [MeSH Terms] OR “Schistosomiasis” [All Fields]) and (2) (“drug name” [MeSH Terms] OR “drug name” [All Fields]) AND (“Schistosomiasis” [MeSH Terms] OR “Schistosomiasis” [All Fields]).

## Results

### Development of a predicted target list

In the TDR Targets Database, 59 genes were identified that encode *S. mansoni* energy metabolism enzymes. All the products of these genes were considered as potential therapeutic targets (Table 3).

### Identification of targets and drugs with potential for anti-schistosomiasis therapy

The DrugBank and TTD databases returned 11 and one targets, respectively, with approved drugs against them. The protein targets and their respective functions are presented in Table 1.

For each of the targets, the DrugBank and TTD databases returned one or more drugs. Twenty drugs were identified, 19 in the DrugBank and one in the TTD. These drugs and their original indications are presented in Fig 2.

**Fig 2.**
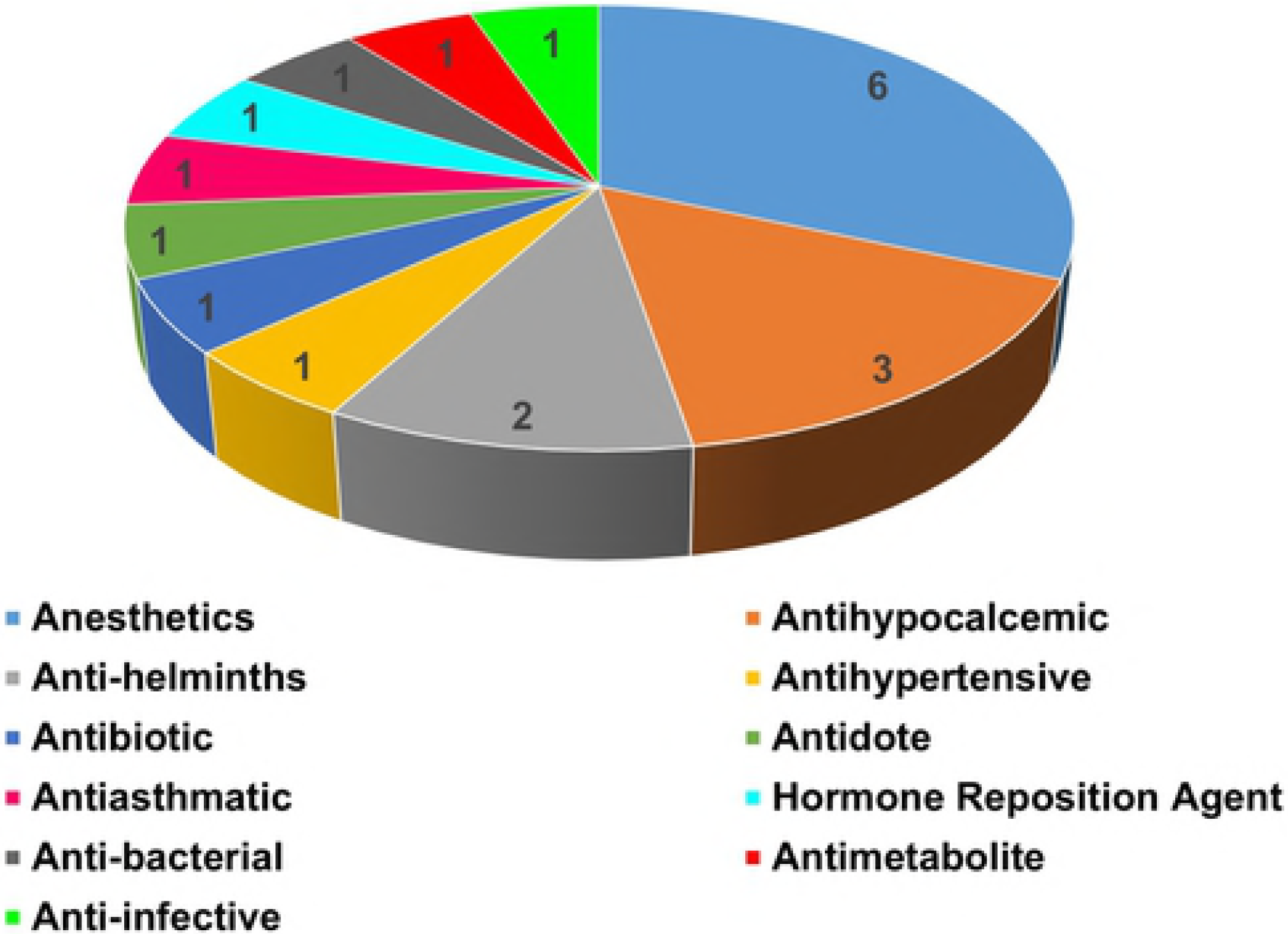
Original principal drug indications with potential anti-schistosomiasis activity.

Of the ten targets, eight had high chemotherapeutic potential because of their very low E-values. Five targets that interact with seven high potential anti-schistosomiasis drugs were selected. Detailed information on these compounds is presented in Table 2.

### Identification of interaction network between drug(s), target(s) protein(s), and side effect(s)

In this analysis, drugs identified as having therapeutic potential against schistosomiasis were considered if they had a low cost, were approved for use in humans, and possessed few side effects. Interactions of drugs identified with other targets were investigated using the Promiscuous database to meet this last criterion. Each of the drugs was subjected to analysis, and 18 of them presented interactions with different molecules. In this study, only the interaction networks of drugs with high anti-schistosomiasis potential were detailed (Table 2).

### Thiabendazole

Thiabendazole shows interaction potential with the targets cytochrome P450 1A2 (*CP1A2_HUMAN*) and cytochrome P450 1A1 (*CP1A1_HUMAN*). Interactions and side effects are presented in Fig S1.

### Albendazole

Albendazole shows interaction potential with the targets cytochrome P450 3A4 (*CP3A4_HUMAN*), cytochrome P450 1A2 (*CP1A2_HUMAN*), and cytochrome P450 1A1 (*CP1A1_HUMAN*). Interactions and side effects are shown in Fig S2.

### Fomepizole

Fomepizole shows interaction potential with the targets alcohol dehydrogenase 1A (*ADH1A_HUMAN*), alcohol dehydrogenase 1B (*ADH1B_HUMAN*), Alcohol dehydrogenase 1C (*ADH1G_HUMAN*), catalase (*CATA_HUMAN*), cytochrome P450 2A6 (*CP2A6_HUMAN*), and cytochrome P450 2E1 (*CP2E1_HUMAN*). Interactions and side effects are shown in Fig S3.

### Desflurane

Desflurane presents interaction potential with targets Glutamate receptor 1 (*GRIA1_HUMAN*), Glycine receptor alpha-1 subunit (*GLRA1_HUMAN*), ATP delta synthase subunit, mitochondrial (*ATPD_HUMAN*), gamma-aminobutyric acid receptor alpha-1 subunit (*GBRA1_HUMAN*), potassium voltage-gated channel subfamily A member 1 (*KCNA1_HUMAN*), and ATPase for calcium transport type 2C member 1 (*AT2C1_HUMAN*). Interactions and side effects are shown in Fig S4.

### Isoflurane

Isoflurane presents interaction potential with targets ATPase member 2C of calcium transport type 1 (*AT2C1_HUMAN*), ATP delta synthase subunit (*ATPD_HUMAN*), calmodulin (*CALM_HUMAN*), cytochrome P450 2E1 (*CP2E1_HUMAN*), cytochrome P450 2B6 (*CP2B6_HUMAN*), gamma-aminobutyric acid receptor alpha-1 subunit (*GBRA1_HUMAN*), Glycine alpha-1 receptor subunit (*GLRA1_HUMAN*), Glutamate 3 receptor (*GRIA3_HUMAN*), Glutamate 1 receptor (*GRIA1_HUMAN*), NADH-ubiquinone oxidoreductase (*NU1M_HUMAN*), and potassium voltage-gated channel for subfamily 1 member 1 (*KCNA1_HUMAN*). Interactions and side effects are shown in Fig S5.

### Alendronate

Alendronate presents interaction potential with the Tyrosine phosphatase type 4 (*PTN4_HUMAN*) and farnesyl pyrophosphate synthase targets (*FPPS_HUMAN*). Interactions and side effects are shown in Fig S6.

### Diazoxide

Diazoxide demonstrates interaction potential with targets ATP-binding cassette subfamily C member 8 (*ABCC8_HUMAN*), ATPase transport of sodium/potassium alpha 1 subunit (*AT1A1_HUMAN*), ATP-sensitive inward rectifier potassium channel 11 (*IRK11_HUMAN*), calcium-activated potassium channel subunit alpha-1 (*KCMA1_HUMAN*), sulfonylurea receptor (*Q59GM5_HUMAN*), carbonic anhydrase 1 (*CAH1_HUMAN*), carbonic anhydrase 2 (*CAH2_HUMAN*), carbonic anhydrase 4 (*CAH4_HUMAN*), and solute carrier family 12 member 3 (*S12A3_HUMAN*). Interactions and side effects are shown in Fig S7.

## Discussion

### Drugs already tested against *Schistosoma* spp. metabolism

Two drugs identified using this strategy were already tested as anti-schistosomiasis agents: thiabendazole and albendazole.

Thiabendazole is indicated for the treatment of parasitic illnesses caused by *Ascaris lumbricoides*, *Strongyloides stercoralis*, *Necator americanus*, *Ancylostoma duodenale*, *Ancylostoma braziliense*, *Trichuris trichiura, Toxocara canis*, *Toxocara cati*, and *Enterobius vermicularis*. Thiabendazole acts as an inhibitor of the flavoprotein subunit of fumarate reductase in these organisms, which catalyzes the oxidation of succinate to fumarate during the Krebs cycle (Drug Bank Data). Here, it is suggested that this drug exhibits potential as a succinate dehydrogenase inhibitor (Table 2), as it interferes with the transfer of electrons from succinate to ubiquinone (*Smp_061230*) (Gene DB Data).

Albendazole is an anthelmintic used to treat parasitic illnesses caused by *Taenia solium* and *Echinococcus granulosus*. This drug acts as a beta tubulin inhibitor and interferes with microtubule polymerization and assembly. As a consequence of cytoplasmic microtubule loss, a reduction in glucose absorption by the parasites in larval and adult stages occurs, resulting in reduced ATP synthesis. Moreover, albendazole acts as a fumarate reductase inhibitor, a flavoprotein subunit present in the metabolism of *Shewanella oneidensis*, interfering with the catalysis of the reduction of fumarate (Drug Bank Data). *In silico* analysis demonstrated that albendazole inhibits succinate dehydrogenase (*Smp_061230*) (Table 2).

To evaluate *in vivo* anti-schistosomiasis activity of thiabendazole and albendazole in mice infected with *S. mansoni*, 100 mg/kg/day and 500 mg/kg/day doses were used, respectively. In this assessment, it was found that albendazole treatment showed no effect on the number of dead adult specimens and/or the elimination of eggs from this parasite. In contrast, treatment using thiabendazole significantly increased the mortality of adult worms [32].

### Drugs not already tested against *Schistosoma* spp

The study’s main purpose was to identify new anti-schistosomiasis drugs using an *in silico* repositioning strategy with drugs approved for treating other diseases. The following is a discussion of each of the drugs identified in the present study and their potential effects on *S. mansoni* energy metabolism.

Fomepizole is indicated for treating methanol poisoning or used when the ingestion of ethylene glycol is suspected [33]. This drug acts as an inhibitor of alcohol dehydrogenase 1B, present in human metabolism [34]. In this target, fomepizole inhibits the oxidation of ethanol to acetaldehyde (DrugBank data). In the present study, fomepizole was identified as having the capacity to inhibit (*Smp_044440*) alcohol dehydrogenase (Table 2), as both cited targets present similar functions. Results obtained from the Promiscuous database reveal that fomepizole also inhibits catalase (*CATA_HUMAN*), which degrades hydrogen peroxide; cytochrome P450 2A6 (*CP2A6_HUMAN*), which functions as a component in the metabolic activation of the B1 aflatoxin; and cytochrome P450 2E1 (*CP2E1_HUMAN*), which inactivates a series of drugs and xenobiotics and also bioactivates many xenobiotic substrates in their hepatotoxic or carcinogenic forms (Fig S3) (Promiscuous data).

Desflurane is used as an inhalation agent for inducing and maintaining general anesthesia (DrugBank data). This drug inhibits the mitochondrial ATP synthase subunit delta target, in human metabolism. This inhibitor interferes with the synthesis of ATP from ADP in the presence of an electrochemical gradient [35]. *In vitro* analysis verified that desflurane presents antiplatelet activity, interfering with the induction of platelet and leukocyte aggregation, through inhibiting the stimulation of P-selectin synthesis, which carries out the function of heterophilic cell adhesion [36]. Desflurane was identified as presenting potential to inhibit mitochondrial ATP synthase delta chain (*Smp_082120*) (Table 2), by interfering with the proton transport through a membrane, generating an electrochemical gradient that contributes to ATP synthesis (Gene DB Data). Moreover, desflurane presents potential target interaction with Glutamate 1 receptor (*GRIA1_HUMAN*), which regulates glutamate release; Glycine subunit alpha-1 receptor (*GLRA1_HUMAN*), which regulates glycine release; NADH-ubiquinone oxidoreductase (*NU1M_HUMAN*), which transfers NADH electrons to the respiratory chain; ATP synthase delta mitochondrial subunit (*ATPD_HUMAN*), which produces ATP from ADP in the presence of a proton gradient; gamma-aminobutyric acid subunit alpha-1 receptor (*GBRA1_HUMAN*), which functions in regulating histamine release; potassium voltage-gated channel subfamily A member 1 (*KCNA1_HUMAN*), which functions in transmembrane potassium transport; and ATPase of calcium transport type 2C member 1 (*AT2C1_HUMAN*), an ATP hydrolysis catalyst that is also involved in calcium transport (Fig S4) (Promiscuous data).

Isoflurane is also used for inducing and maintaining general anesthesia. This drug acts as an inhibitor of the mitochondrial ATP synthase subunit delta target, in human metabolism. This inhibitor interferes with the production of ATP from ADP in the presence of a proton gradient (DrugBank data). *In vivo* analysis verified that isoflurane presents potential as an antidepressant, inhibiting the glycogen synthase kinase 3β target, interfering with the regulatory functions of hormonal homeostasis and Wnt signaling (pathway associated with cell proliferation) [37]. This study identified that isoflurane can inhibit mitochondrial ATP synthase subunit delta (*Smp_082120*) (Table 2), as both targets mentioned have similar functions.

Even though desflurane and isoflurane offer anesthetic pharmacological activity, it is possible to make them usable for clinical trials against *S. mansoni*, using molecular remodeling.

Alendronate is indicated for treatment of Paget’s disease and osteoporosis. This drug acts as an inhibitor of the ATPase type IV subunit A target, interfering in proton transport through a rotation mechanism. Alendronate exhibits toxicity evidenced by esophageal mucosa damage (DrugBank data). In this study, alendronate was identified as having the capacity to inhibit ATP synthase subunit beta (*Smp_038100*), interfering with the function of proton transport connected to ATP synthesis (Table 2) (Gene DB data). Furthermore, alendronate presents interaction potential with the Tyrosine phosphatase type 4 target (*PTN4_HUMAN*), which has roles in the junctions between the membrane and the cytoskeleton and with the farnesyl pyrophosphate synthase target (*FPPS_HUMAN*), through the function of the biosynthesis of isoprenoids that catalyze the formation of farnesyl diphosphate (Fig S6) (Promiscuous data).

## Conclusions

This study presents an *in silico* repositioning strategy to identify new drugs for treating intestinal schistosomiasis based on the principle of target similarity to identify drugs approved for clinical use in humans. Twenty drugs were identified, eight of which have high schistosomicide potential. The action of the drugs was compared among the targets using databases that presented interaction analysis of the drug with other targets. Using this methodology, it was possible to confirm the drug-protein target interaction, as well as the interaction with other targets that may or may not have roles in energy metabolism. Furthermore, information was obtained on possible side effects of the identified drugs. However, it is important to emphasize that there are no guarantees that the identified compounds will act on *S. mansoni* energy metabolism targets.

## Financial resources

This work was supported by the Brazilian National Council for Scientific and Technological Development (CNPq/MS/DECIT n. 37/2014. Projeto 470298/2014-6).

